# Structural and mutational analyses define distinct molecular routes to broad SARS-CoV-2 receptor-binding domain recognition

**DOI:** 10.64898/2026.08.17.745277

**Authors:** Morgan E. Abernathy, William B. Foreman, Jasmyn A. Lopez, Viren A. Baharani, Danielle Vahdat, Yu E. Lee, Michael R. Eso, Zijun Wang, Paul D. Bieniasz, Michel C. Nussenzweig, Tyler N. Starr, Christopher O. Barnes

## Abstract

Broadly reactive antibodies elicited by SARS-CoV-2 infection or vaccination can reveal conserved viral vulnerabilities and inform vaccines with broad coronavirus coverage. Here, we characterize two human-derived monoclonal antibodies, B2014 and C5078, that recognize conserved epitopes on the SARS-CoV-2 RBD and retain activity across antigenically distinct variants. Notably, C5078 also recognizes diverse sarbecoviruses and remains active against currently circulating variants, including XFG and NB.1.8.1. Cryo-EM structures reveal that B2014 recognizes an epitope adjacent to the class 3 antibody site, whereas C5078 targets the highly conserved, cryptic site V epitope. Structural analysis defines how C5078 uses affinity-matured interactions to engage conserved RBD residues, providing a molecular basis for its exceptional breadth. Deep mutational scanning across multiple SARS-CoV-2 variant backgrounds further defines potential pathways of antibody escape, explaining the loss of B2014 activity against antigenically evolved variants while revealing a high barrier to escape from C5078. Together, these findings define distinct structural solutions for broad RBD recognition and highlight conserved, mutationally constrained epitopes that may serve as targets for vaccines designed to elicit antibody responses resilient to ongoing SARS-CoV-2 evolution and future sarbecovirus emergence.

## INTRODUCTION

Since its emergence in 2019, SARS-CoV-2 has undergone extensive antigenic evolution, repeatedly diminishing the effectiveness of antibody responses elicited by prior infection, vaccination, and therapeutic monoclonal antibodies (mAbs). Viral entry is mediated by the trimeric spike glycoprotein, whose receptor-binding domain (RBD) engages the host receptor angiotensin-converting enzyme 2 (ACE2) to initiate the conformational changes required for membrane fusion (1, 2). Because the RBD is a dominant target of neutralizing antibodies, many of the earliest and most potent SARS-CoV-2 mAbs recognized epitopes overlapping the ACE2-binding site (3, 4). However, the antigenic plasticity of these immunodominant surfaces has enabled successive viral variants to escape most early RBD-directed antibodies, including those developed for clinical use (5–7). The continued emergence of antigenically divergent variants, together with the threat of future zoonotic sarbecovirus spillovers, has therefore intensified efforts to identify conserved sites of antibody vulnerability that are less permissive to viral escape (8).

Despite substantial sequence diversity within the RBD, several conserved surfaces can support broad antibody recognition. Whereas class 1 and 2 antibodies predominantly target the highly variable ACE2 receptor-binding motif, antibodies directed toward more conserved surfaces outside this region can retain activity across antigenically divergent SARS-CoV-2 variants and, in some cases, diverse sarbecoviruses (9–11). These include class 3 antibodies as well as antibodies targeting cryptic epitopes that are poorly accessible within the closed prefusion spike. Among the latter, class 5, or site V, antibodies recognize a highly conserved surface that becomes accessible through conformational opening or rearrangement of the RBD, whereas class 6 antibodies recognize an adjacent region spanning the class 3 and site V surfaces (10–13). Together, these antibodies demonstrate that the RBD contains conserved vulnerabilities capable of supporting broad neutralization despite extensive viral antigenic evolution.

Site V, in particular, has emerged as a particularly promising target for broad and escape-resistant antibody responses. Antibodies targeting this cryptic surface can recognize diverse SARS-CoV-2 variants and sarbecoviruses, consistent with structural and functional constraints that limit variation within the epitope. Recent studies have revealed diverse immunogenetic and structural solutions for site V recognition and suggest that engagement of this normally occluded surface can promote substantial conformational rearrangement or destabilization of the spike trimer (11, 13). However, the molecular interactions that enable individual antibodies to exploit this constrained surface, the consequences of site V engagement for intact spike architecture, and the extent to which sequence conservation translates into resistance to prospective antibody escape remain incompletely defined. Moreover, although antibodies targeting conserved cryptic RBD epitopes have demonstrated protective activity in vivo, the protective potential of site V-directed antibodies remains less well established.

Here, we characterize two human RBD-directed antibodies, B2014 and C5078, that exemplify distinct routes to breadth. B2014 recognizes a conserved epitope adjacent to the class 3 site and retains activity against multiple antigenically divergent SARS-CoV-2 variants, whereas C5078 targets site V, recognizes RBDs from diverse sarbecoviruses, and remains active against recently circulating SARS-CoV-2 variants. Using cryo-electron microscopy, we define the molecular basis of recognition by both antibodies and show how affinity-matured interactions enable C5078 to engage highly conserved residues within site V. Analysis of C5078 bound to intact spike further reveals a highly splayed RBD configuration, providing structural evidence linking site V engagement to substantial spike rearrangement. We integrate these structures with deep mutational scanning across multiple SARS-CoV-2 variant backgrounds to define potential pathways of escape and demonstrate the limited capacity of single RBD substitutions to disrupt C5078 recognition. Finally, we evaluate the protective potential of C5078 following viral challenge in vivo. Together, these findings define distinct molecular routes to broad RBD recognition and establish site V as a conserved, mutationally constrained vulnerability with a high barrier to antibody escape.

## RESULTS

### RBD-specific mAbs B2014 and C5078 exhibit distinct patterns of breadth across SARS-CoV-2 variants and diverse sarbecoviruses

We previously identified two broadly reactive RBD-directed monoclonal antibodies, B2014 (IGHV1-46/IGLV1-40) and C5078 (IGHV5-51/IGLV3-21), from independent donor cohorts (14, 15). B2014 was isolated from a vaccinated donor following Omicron breakthrough infection, whereas C5078 was identified from an elderly individual following a third mRNA vaccine dose with no history of SARS-CoV-2 infection. At the time of their discovery, B2014 exhibited broad activity against circulating SARS-CoV-2 variants with a limited mutational escape profile (16), while C5078 additionally recognized diverse sarbecoviruses, motivating further structural and functional characterization to determine the molecular basis and evolutionary durability of their breadth.

To understand the binding properties of B2014 and C5078, we first measured Fab binding kinetics to Wuhan-Hu-1 RBD by biolayer interferometry (BLI). Both antibodies exhibited strong binding affinities to Wuhan-Hu-1 RBD with slow dissociation rates (Fig. 1A). We next assessed antibody resilience to SARS-CoV-2 antigenic evolution by measuring IgG binding to RBDs derived from a panel of SARS-CoV-2 variants of concern (VOCs) by Enzyme-Linked Immunosorbent Assay (ELISA). Of the tested strains, B2014 maintained binding across early variants and multiple Omicron lineages but lost detectable binding beginning with BA.2.86 (Fig. 1B). In contrast, C5078 retained binding to all RBDs tested, including the recently circulating XFG and NB.1.8.1 variants (Fig. 1B). Consistent with these binding profiles, pseudovirus neutralization assays demonstrated that B2014 progressively lost neutralizing activity against antigenically evolved VOCs, whereas C5078 retained neutralizing activity, with a profile comparable to the broadly neutralizing site V antibody S2H97 (Fig. S1). Across susceptible SARS-CoV-2 variants, both antibodies retained potent neutralizing activity, with IC50 values ranging from 0.15-0.8 µg/mL for C5078 and 0.01-0.05 µg/mL for B2014, consistent with their previously reported neutralization potencies against early SARS-CoV-2 VOCs (14, 16). Given the resilience of C5078 to extensive SARS-CoV-2 antigenic evolution, we next assessed its recognition of RBDs from diverse sarbecoviruses using a multiplexed yeast-display library spanning five phylogenetic clades (Fig. 1C, Fig S2A). C5078 exhibited broad cross-reactivity, recognizing RBDs across multiple strains within clades 1b, 2, and 3, with binding profiles that overlapped substantially with those of the pan-sarbecovirus site V antibody S2H97 (Fig. 1C). These results indicate that C5078 recognizes an RBD surface that is conserved not only across SARS-CoV-2 VOCs but also among phylogenetically diverse sarbecoviruses.

**Figure 1.**
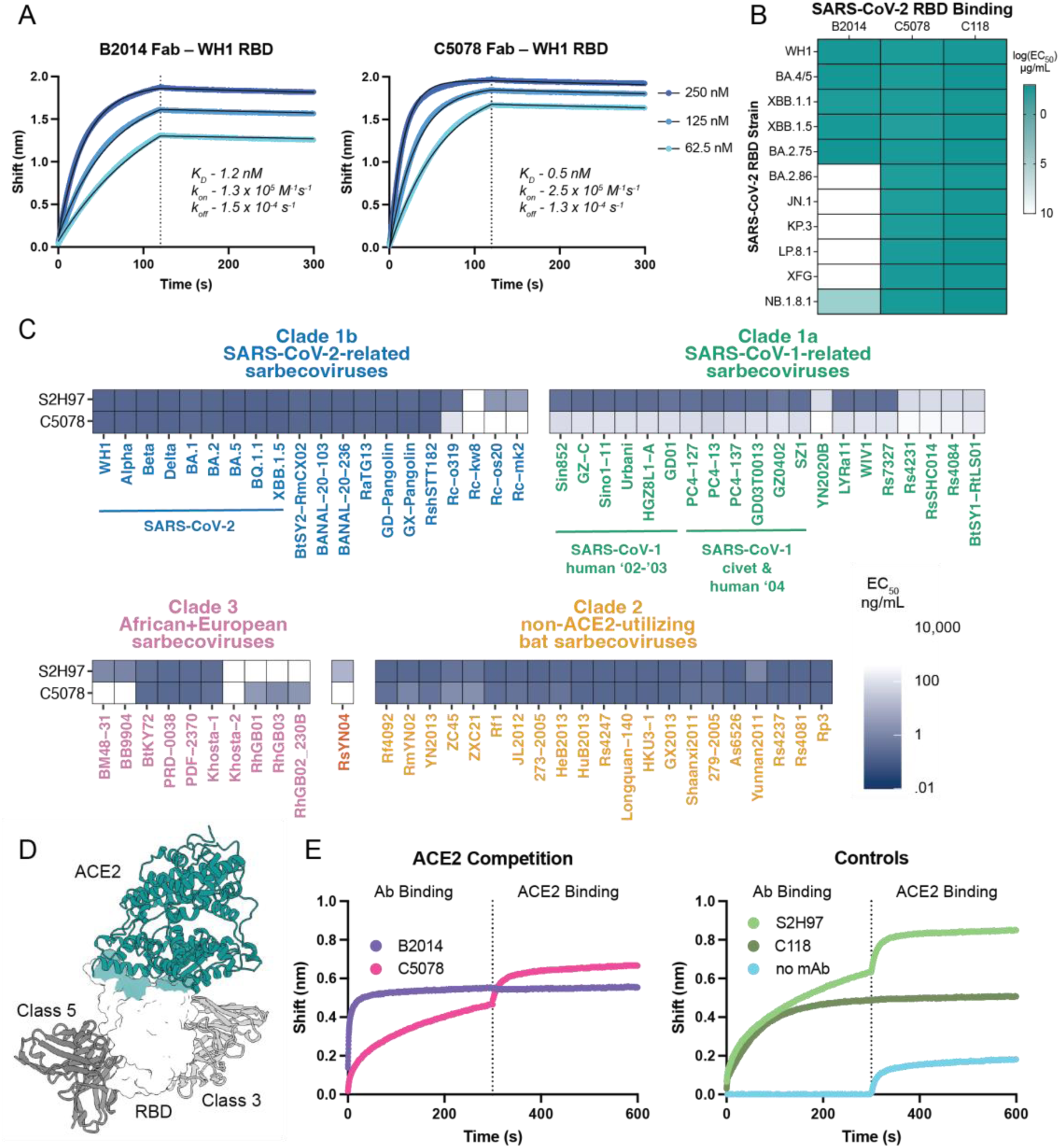
RBD-specific mAbs B2014 and C5078 exhibit broad recognition of SARS-CoV-2 variants and diverse sarbecoviruses. **A)** Biolayer interferometry (BLI) association and dissociation curves for B2014 and C5078 Fabs binding to biotinylated Wuhan-Hu-1 (WH1) RBD immobilized on streptavidin biosensors at the indicated Fab concentrations. Dissociation begins at 120 s (dotted line). Fits are shown as black solid lines, with derived kinetic parameters indicated. **B)** Heatmap of ELISA log(EC50) values of monoclonal antibodies binding to RBDs from the indicated SARS-CoV-2 variants. **C)** Pan-sarbecovirus RBD binding for mAbs S2H97 and C5078. Each antibody was measured for binding against each sarbecovirus RBD using a multiplexed FACS-seq titration assay with a yeast-display library of RBDs. Color-coded EC50 values represent the geometric mean across internally replicated barcodes within the library linked to the same RBD. **D)** Model of SARS-CoV-2 RBD bound to its receptor human angiotensin-converting enzyme 2 (ACE2) (PDB 6LZG) with representative antibodies (aligned by bound RBD) from Class 3 (S309 Fab; PDB 7XSW) and Class 5 (S2H97 Fab; PDB 7M7W) shown in shades of gray, and ACE2 epitope as teal surface. **E)** Competition BLI assay assessing whether B2014 and C5078 interfere with ACE2 binding to SARS-CoV-2 RBD. Antibodies were allowed to associate for 300 s, followed by a 300 s ACE2 association phase. S2H97 and 88 were included as noncompeting and competing controls, respectively.

Given their distinct patterns of breadth and neutralization, we next sought to determine whether B2014 and C5078 use different mechanisms to inhibit viral entry. Prior competition-based epitope mapping classified B2014 as a class 3 antibody and C5078 as a site V antibody, indicating recognition of epitopes outside the ACE2 binding site (Fig. 1D). We therefore tested whether either antibody could nevertheless interfere with ACE2 engagement using a BLI-based competition assay. Binding of C5078 with the RBD had no detectable effect on subsequent ACE2 binding, consistent with its assignment to site V and with the behavior of the control site V mAb S2H97 (Fig. 1E). In contrast, B2014 inhibited ACE2 binding despite its previous classification as a class 3 antibody. This unexpected competition suggests that the B2014 binding footprint or orientation sterically interferes with receptor engagement, providing a potential mechanism of neutralization. These distinct functional properties prompted us to determine the structural basis of RBD recognition by B2014 and C5078.

### B2014 recognizes a class 3-adjacent epitope vulnerable to viral escape

To define the structural basis of B2014 recognition, we complexed excess B2014 Fab with BA.1 S6P (**Fig. S3A**) and determined a single particle cryo-EM map to 3.0 Å (**Fig. S4A**). A locally refined reconstruction of one Fab-RBD interface was resolved to 3.4 Å resolution and was used for model building and analysis (**Fig. 2A; Fig. S4A**). B2014 recognizes an epitope adjacent to the canonical class 3 site, adopting a distinct binding orientation compared with representative class 3 mAbs, including C135 and S309 (**Fig. S3B**). Its footprint extends toward the ACE2-binding site, providing a structural explanation for its ability to directly compete with ACE2 binding as observed by BLI (**Fig. 1E**; **Fig. 2B**).

**Figure 2.**
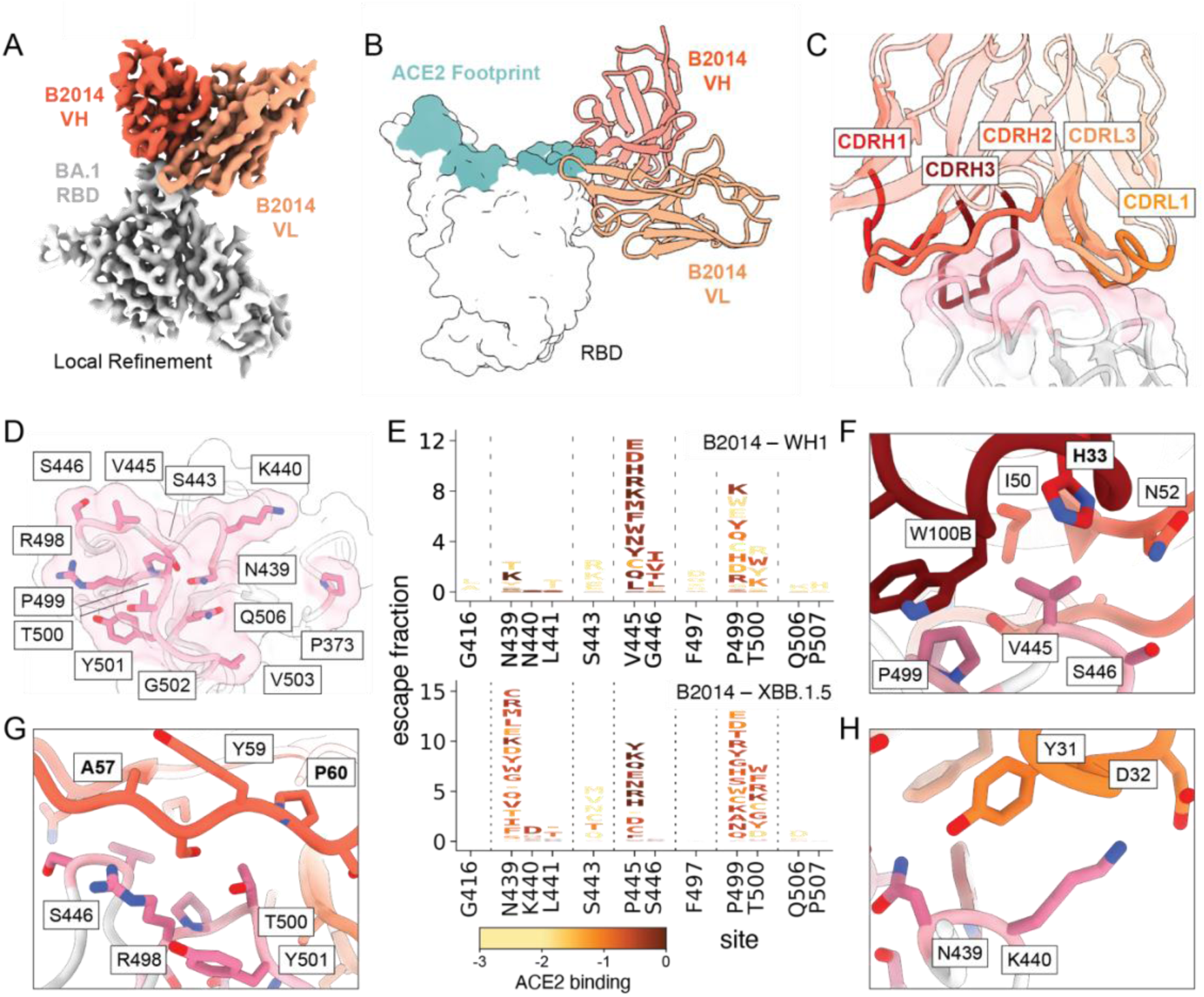
B2014 binds adjacent to the Class 3 site on the SARS-CoV-2 RBD. **A)** 3.4 Å cryo-EM local map reconstruction of the B2014 VHVL–BA.1 RBD complex. The B2014 heavy and light chains are shown in orange and light chain, respectively, with RBD shown in gray. This color scheme is maintained throughout the figure. **B)** Binding orientation of B2014 relative to the ACE2-binding footprint on the SARS-CoV-2 RBD (shown in the same orientation as Fig. 1D). **C)** B2014 paratope comprising CDR loops in shades of orange and RBD epitope (pink). **D)** B2014 epitope on the BA.1 RBD, with all contacting residues shown as sticks. **E)** Escape maps for B2014 in SARS-CoV-2 strains Wuhan-Hu-1 and XBB.1.5 from deep mutational scanning. Escape mutations are visualized in logoplots, where the height of each letter indicates the strength of escape from antibody binding conferred by each mutation. Mutations are colored by their separately measured impact on ACE2-binding affinity (28, 39). See Fig. S2 for additional detail. **F-H)** Molecular interactions between B2014 and the RBD highlighting **(F)** RBD residues V445 and S446, which were identified by DMS as critical contacts and are mutated in BA.2.86; **(G)** commonly mutated residues within the RBD loop spanning residues 498-501 and their interactions with CDRH2; and **(H**) commonly mutated RBD residues N439 and K440 and their interactions with CDRH1. Antibody residues are colored as in (**C**), with somatic hypermutations (SHMs) indicated in bold.

B2014 utilizes five of six CDR loops for antigenic recognition and engages the RBD loops spanning residues 439-446, 498-506, and P373 with a total buried surface area (BSA) of 740 Å2 (**Fig. 2C,D; Fig. S3C,D**). To determine how mutations within and surrounding the RBD footprint affect antibody recognition, we performed deep mutational scanning (DMS) to measure the impacts of virtually all single amino acid mutations in the Wuhan-Hu-1 and XBB.1.5 RBD backgrounds (**Fig. 2E; Fig S2**). Escape was concentrated at residues within the structurally defined B2014 epitope, including positions 439 and 445, which have undergone substitutions in earlier variants (17). Notably, substitutions at 445 strongly disrupted B2014 recognition, and BLI confirmed substantially reducing binding to RBDs containing V445E or V44L (**Fig. S5A**). Consistent with these findings, the V445H substitution emerged in BA.2.86 (18), the first variant in our panel against which B2014 lost detectable binding (**Fig. 1B; Fig. S5C**).

The B2014-BA.1 RBD structure provides a molecular explanation for the sensitivity of B2014 to variation at this position. V445 lies within a predominantly hydrophobic antibody-antigen interface formed by B2014 residues H33, I50, and W100B. Thus, substitutions at this site with charged or bulky amino acids results in disruption of this interface, consistent with the strong escape profiles shown by such amino acids by DMS (**Fig 2E,F**). Supporting the importance of this site, among a previously described panel of synthetic SARS-CoV-2 variants, all variants resistant to B2014 contained the V445E substitution, whereas the remaining substitutions present in these variants were also found in B2014-sensitive backgrounds (16). Furthermore, V445H was the only substitution within the structurally defined B2014 epitope that distinguished the B2014-sensitive strain BA.2.75 from the resistant BA.2.86 RBD. Together with our DMS and BLI validation, these independent observations identify residue 445 as a key determinant of B2014 escape.

Beyond V445, B2014 contacts the adjacent RBD loop spanning residues 498–501 primarily through CDRH2, while residues N439 and K440 are positioned to interact with CDRH1 (**Fig. 2G,H**). These sites also exhibit sensitivity to mutation in our DMS analysis, indicating additional potential pathways of escape. Thus, although B2014 tolerated substantial early SARS-CoV-2 antigenic evolution, its reliance on several mutable RBD residues ultimately created vulnerabilities exploited by later variants. These findings provide a useful contrast to C5078, prompting us to investigate whether its sustained activity against contemporary variants reflects recognition of a more conserved and mutationally constrained RBD surface.

### C5078 binds the conserved site V cryptic epitope and promotes an open spike conformation

To define the structural basis for the breadth of C5078, we determined a 2.4 Å cryo-EM structure of C5078 Fab bound to the SARS-CoV-2 RBD in a complex containing S2X259 Fab (19) and S309 Fab (20) to increase complex size and facilitate structure determination (**Fig. 3A**, **Fig 4B**). A 2.3 Å locally refined reconstruction focused on the C5078 V_H_V_L_ – RBD complex was used for model building and analysis of antibody-antigen contacts (**Fig. S4B**). The structure confirms that C5078 recognizes the cryptic site V epitope, distal from the ACE2 binding site and largely occluded by the neighboring N-terminal domain (NTD) when the RBD adopts the “down” conformation (**Fig 3B**). Given these interdomain Spike interactions, site V is among the most conserved surfaces of the RBD, with mutations within this region negatively affecting RBD folding and expression (11–13, 21).

**Figure 3.**
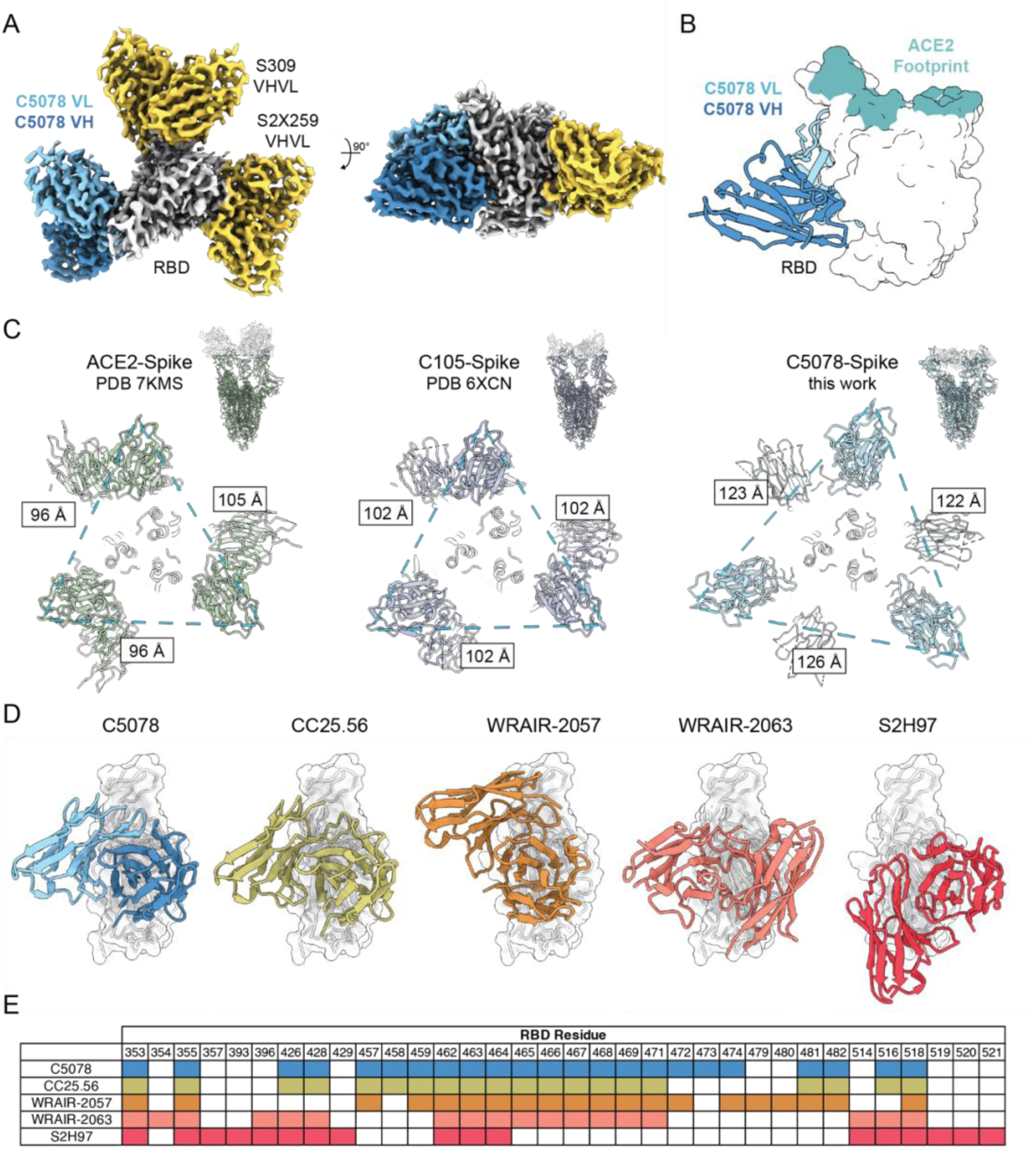
C5078 targets the conserved site V cryptic epitope on the SARS-CoV-2 RBD. **A)** 2.3 Å cryo-EM global map of the C5078–RBD–S309–S2X259 complex. The C5078 heavy and light chains are shown in dark blue and sky blue, respectively, and the RBD is shown in gray. Additional Fabs are shown yellow. This color scheme is maintained throughout. **B)** Binding orientation of C5078 relative to the ACE2-binding footprint on the SARS-CoV-2 RBD (shown in the same orientation as Fig. 1D). **C)** Comparison of inter-protomer RBD distance computed for the “Up” RBDs in ACE2-bound Spike (left), Class 1 C105-bound Spike (middle), and C5078-bound Spike (right). For C5078, three RBD subunits and three Spike protomers missing the RBD subunit were each fit separately into the low-resolution map from Fig S6B before distances were calculated. Distances calculated between G446 Cα atoms of each RBD using ChimeraX. **D)** Comparison of the C5078 binding pose with previously characterized site V antibodies. Fab-RBD structures were superimposed by alignment of the RBDs and are shown in the same orientation. From left to right: C5078 (this work, PDB 37JA), WRAIR-2057 (PDB 7N4I), WRAIR-2063 (PDB 8EOO), and S2H97 (PDB 9ATM). **E)** Comparison of antibody-contacting RBD residues by the indicated site V RBD antibodies. Contacts were defined using a 4.1 Å interchain distance cutoff in ChimeraX.

**Figure 4.**
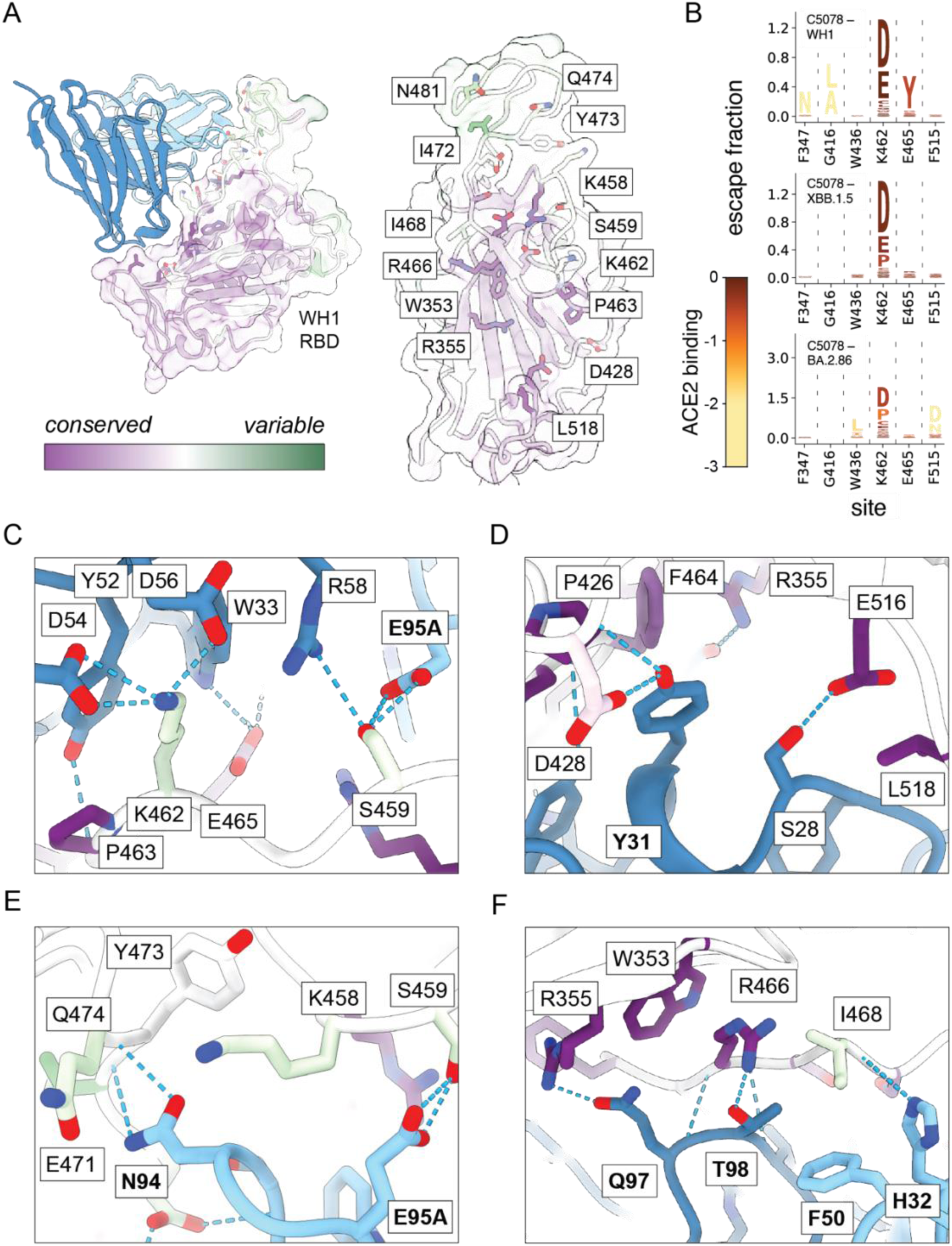
Molecular basis of C5078 binding breadth across sarbecoviruses. **A)** Surface representation of the SARS-CoV-2 RBD colored according to residue-level conservation across sarbecoviruses, as calculated using ConSurf (Ref). The C5078-bound RBD was aligned to the displayed RBD (PDB 7RKU), with side chains of C5078-contacting residues shown. Right, a detailed view of the C5078 epitope with selected contacting residues labeled. **B)** Escape maps for C5078 produced using deep mutational scanning; details as in Fig. 2E. **C-F)** Molecular interactions between C5078 and the RBD highlighting: **C**) selected residues identified as sites of escape by DMS, including CDRH2-mediated contacts; **D**) interactions mediated by CDRH1; **E**) interactions mediated by the SHMs in CDRL3; **F**) interactions mediated by SHMs in CDRH3, CDRL1, and CDRL2 loops. Antibody residues are colored by chain with somatic hypermutations (SHMs) indicated in bold. RBD residues are colored by residue-level conservation as in (A), ranging from highly conserved (purple) to variable (green). Potential electrostatic interactions are shown as cyan dashed lines.

Site V accessibility within the context of the intact spike presents an apparent structural constraint on antibody recognition. Given that previous studies have suggested that site V targeting antibodies can promote spike opening or destabilization of the trimeric Spike (11, 13, 22, 23), we sought to determine how C5078 engages the trimeric spike. C5078 Fab bound prefusion-stabilized spike trimer by BLI (**Fig. S6A**), and single-particle cryo-EM revealed a low-resolution reconstruction of C5078 bound to the BA.1 S6P glycoprotein (**Fig. S6B**). Although the resolution precluded atomic modeling, the reconstruction accommodated three independently fitted spike protomers and three copies of Fab bound to an “up” RBD subunit (based on the high-resolution C5078-RBD model), revealing all three RBDs in an extensively open configuration (**Fig. 3C; Fig. S6B**).

Notably, the C5078-bound RBDs were more widely separated than those observed in representative ACE2- or class 1 antibody-bound open spike structures (**Fig. 3C**). This highly splayed configuration provides structural evidence that simultaneous engagement of the cryptic site V epitope requires, or stabilizes, substantial rearrangement of the spike trimer. These observations complement previous negative-stain EM and biochemical studies suggesting that site V antibodies can destabilize spike and promote S1 shedding. Although the limited resolution prevents us from establishing whether C5078 induces this conformation or preferentially captures a preexisting state, the pronounced opening provides a structural framework for understanding how site V antibodies access an epitope that is otherwise occluded in the closed spike. These observations further suggest that the restricted accessibility of site V may help explain why many antibodies targeting this highly conserved epitope exhibit relatively weak or non-neutralizing activity despite recognizing one of the most evolutionarily constrained surfaces on the RBD.

Nevertheless, multiple antibodies have independently evolved solutions to engage this cryptic surface. Comparison with previously characterized site V antibodies revealed that C5078 most closely resembles CC25.56 (13) in overall pose and number of shared contacts, likely due to usage of the same *IGHV5-51/IGLV3-21* germline genes (**Fig. 3D,E; Fig. S6C,D**). Interestingly, other site V mAbs, including WRAIR-2057 (24) and S2H97 (11), also utilize IGHV5-51 but adopt distinct binding orientations or engage partially overlapping footprints, underscoring the diversity of molecular solutions capable of targeting this cryptic epitope (**Fig. 3D,E**).

### C5078 targets highly conserved and mutationally constrained RBD residues with a high barrier to escape

To better understand the molecular basis for C5078 breadth, we analyzed the specific contacts between C5078 and the RBD in the context of sarbecovirus sequence conservation and SARS-CoV-2 antigenic evolution. The majority of the C5078 epitope is composed of residues that are highly conserved across sarbecoviruses, and nearly all contacting residues remain unmutated in the currently circulating XFG variant (Fig. 4A, Fig. S5D). C5078 engages this surface extensively, burying 1052 Å^2^ of RBD surface area through contacts involving all six complementarity-determining region (CDR) loops (Fig. S6C,D).

We next asked whether conservation of the C5078 epitope translates into resistance to prospective antibody escape. Deep mutational scanning revealed a remarkably restricted escape landscape, with relatively few single amino acid substitutions conferring substantial escape (Fig. 4B). Importantly, this restricted landscape was maintained across antigenically divergent RBD backgrounds, suggesting that the high barrier to C5078 escape is not restricted to the ancestral Wuhan-Hu-1 sequence context. The most consistent escape site across all strains tested was K462, with charge-reversing substitutions exhibiting particularly strong effects (Fig. 4B). Introduction of K462D into the XBB.1.5 RBD increased C5078 dissociation rates by BLI, validating this residue as a potential pathway for escape (Fig. 5B). Despite its strong effect on C5078 recognition, substitutions at K462 have not become prevalent during SARS-CoV-2 evolution, consistent with the continued activity of C5078 against contemporary variants.

The high-resolution structure provides a molecular explanation for both the sensitivity of C5078 to substitution at K462 and its broader resistance to escape. K462 participates in electrostatic interactions with two negatively charged aspartic acid residues in the CDRH2 loop of C5078, D54 and D56, explaining the strong effects of charge-switching substitutions at this position (Fig. 4C). Other DMS-identified sites, including S459 and E465, contribute to a broader interaction network of residues across CDRH1, CDRH2 and CDRL3 (Fig. 4C). However, much of the C5078 footprint is centered on highly conserved residues, including P426, F464, E516, and L518 (Fig. 4D–F), with interactions mediated by CDRH1 residues S28 and Y31, the latter of which resulted from somatic hypermutation. In the C5078 CDRL3 loop, SHM-derived residues N94 and E95A contact the periphery of the epitope closer to the ACE2 binding site (Fig. 4E). Finally, four SHM-derived residues from the CDRL1, CDRL2, and CDRH3 loops contact residues on the edge of the epitope, closer to that of Class 6 binders, which bridge Class 5 and Class 3 (Fig. 4F). Thus, although individual substitutions can disrupt key interactions, the extensive engagement of a conserved and functionally constrained RBD surface limits the number of accessible pathways for single-mutation escape.

### C5078 shows a trend toward reduced viral burden following SARS-CoV-2 challenge

Given the breadth and escape resistance of C5078 *in vitro*, we next evaluated its activity *in vivo* using K18-hACE2 mice challenged with SARS-CoV-2 XBB.1.5. C5078 was administered either 24 h before infection as prophylaxis or 4 h after infection as treatment, and viral burden was subsequently quantified by SARS-CoV-2 RNA levels (Fig. S7). C5078-treated animals exhibited lower mean viral RNA levels in both experimental settings, although these differences did not reach statistical significance. The variability observed among individual animals, including a control animal with relatively low viral RNA levels, reduced the ability to detect statistically significant differences in this cohort. Thus, these results suggest a trend toward antiviral activity *in vivo* but will require additional studies to establish the protective efficacy of C5078 and other site V-directed antibodies.

Collectively, these data establish C5078 as an independently elicited site V antibody whose breadth is supported by extensive recognition of conserved RBD residues, affinity-matured interactions, and a restricted mutational escape landscape that persists across divergent SARS-CoV-2 backgrounds. Together with structural evidence of a highly splayed C5078-bound spike conformation, these results reinforce site V as a conserved RBD vulnerability and provide a mechanistic framework for understanding how antibodies targeting this surface can remain resilient to continued sarbecovirus evolution.

## DISCUSSION

The continued antigenic evolution of SARS-CoV-2 has distinguished antibodies that exhibit breadth against existing variants from those targeting epitopes intrinsically resistant to viral escape. Here, we structurally and functionally characterized two broadly reactive RBD antibodies, B2014 and C5078, that illustrate these distinct outcomes. Although both retained activity through substantial early SARS-CoV-2 evolution, B2014 was ultimately escaped by BA.2.86, whereas C5078 remains active against recently circulating variants and recognizes RBDs from diverse sarbecoviruses. By integrating structural analysis with deep mutational scanning (DMS), our findings identify features underlying these divergent trajectories and highlight epitope constraint as an important determinant of durable antibody breadth.

B2014 recognizes an epitope adjacent to the canonical class 3 site and sterically interferes with ACE2 engagement. DMS identified V445 as a dominant site of escape, consistent with reduced binding to V445 mutants and previous analyses of synthetic escape variants. Notably, V445H emerged in BA.2.86 and is the only substitution within the B2014 footprint between the sensitive BA.2.75 and resistant BA.2.86 variants. Together, these observations illustrate how an antibody can retain activity across numerous variants yet remain vulnerable when viral evolution reaches a permissive escape site. Thus, breadth measured against circulating viruses does not necessarily predict resilience to future antigenic change.

In contrast, C5078 targets site V, a cryptic and highly conserved RBD surface shared across diverse sarbecoviruses. Recent studies have established site V as a target of broadly neutralizing antibodies with diverse immunogenetic origins and binding modes. C5078 most closely resembles CC25.56, including shared *IGHV5-51* and *IGLV3-21* germline usage, providing an independently elicited example of a convergent structural solution for recognizing this surface. Our findings extend these studies by linking the molecular basis of site V recognition to prospective escape across multiple SARS-CoV-2 backgrounds. C5078 engages an extensive RBD surface through all six CDR loops, including affinity-matured contacts with highly conserved residues, while DMS revealed few pathways for single-amino-acid escape. The persistence of this restricted escape profile across divergent RBD backgrounds is notable given that epistasis can alter the effects of antibody escape mutations as SARS-CoV-2 evolves.

K462 was the most consistent site of C5078 escape identified by DMS, and K462D substantially impaired binding in the XBB.1.5 background. The structure shows that K462 participates in an electrostatic interaction network with C5078 CDRH2, explaining the sensitivity to charge-reversing substitutions at this position. Yet K462 has remained largely conserved during SARS-CoV-2 evolution, suggesting that this experimentally accessible escape pathway may face constraints in circulating viruses. Interestingly, naturally occurring substitutions at this position are present in several more divergent sarbecoviruses that exhibit reduced C5078 binding. Although these substitutions were not identified as dominant escape mutations in our SARS-CoV-2 DMS experiments, they may become sufficient to diminish binding when combined with additional sequence divergence elsewhere in the RBD (16). This distinction highlights that the determinants of prospective escape within SARS-CoV-2 are not necessarily identical to those limiting breadth across the broader sarbecovirus lineage. This contrasts with V445, where substitutions compatible with viral evolution ultimately enabled B2014 escape, and illustrates the importance of considering both the effects of mutations on antibody recognition and their evolutionary accessibility. Together, these findings suggest that the structural constraints underlying site V recognition remain largely preserved despite substantial antigenic changes elsewhere in the RBD.

Our structural analyses also provide insight into site V recognition in the context of intact spike. A low-resolution cryo-EM reconstruction revealed three C5078 Fabs bound to RBDs in a highly splayed configuration, with greater separation than observed in representative ACE2- or class 1 antibody-bound open spikes. Previous studies have proposed that site V antibodies can promote spike destabilization and S1 shedding, and the C5078-bound structure provides complementary evidence linking engagement of this cryptic surface to substantial spike rearrangement. Although we cannot determine whether C5078 induces this conformation or captures a preexisting state, these observations suggest that site V-directed neutralization may involve conformational effects beyond direct receptor competition. Lastly, C5078 administration before or shortly after XBB.1.5 challenge was also associated with lower mean viral RNA levels *in vivo*, although these differences did not reach statistical significance. These data therefore do not establish protective efficacy but motivate further investigation of the antiviral potential of C5078 and related site V antibodies in appropriately powered studies.

Together, our findings illustrate how antibodies with similar apparent breadth can differ substantially in their resilience to continued viral evolution. B2014 remained dependent on residues that eventually permitted escape, whereas C5078 targets a conserved RBD surface with a restricted mutational escape landscape across multiple variant backgrounds. By linking structural recognition, viral evolution, and prospective escape, this study reinforces site V as a conserved sarbecovirus vulnerability and suggests that vaccine strategies should prioritize not only broad antibody targets, but epitopes whose functional and structural constraints limit future routes of escape.

## MATERIALS AND METHODS

### Cell Lines

Human embryonic kidney (HEK) 293T cells (*Homo sapiens*, embryonic kidney cells) were used for pseudovirus generation and cultured in Dulbecco’s Modified Eagle Medium (DMEM) supplemented with 10% fetal bovine serum (FBS). HEK293T cells overexpressing human angiotensin-converting enzyme 2 (ACE2; HEK293T_ACE2_) used for pseudovirus titration and neutralization assays were generated as previously described (Ref-bianchini) and were cultured in DMEM supplemented with 10% FBS, 1% non-essential amino acids, 1 mM sodium pyruvate, 1x penicillin/streptomycin, and 5 µg mL^-1^ blasticidin. Expi293F™ cells (Thermo Fisher Scientific, cat. no. A14527) were used for recombinant expression of monoclonal IgGs and viral antigens and were cultured in Expi293 expression medium (Thermo Fisher Scientific, cat. no. A1435014) and according to the vendor’s instructions.

### SARS-CoV-2 pseudotyped reporter viruses

The generation of S-pseudotyped lentivirus was performed as previously described (Ref-Bianchini). In brief, plasmids encoding for a C-terminally truncated spike protein with the transmembrane and cytosolic tail removed (pSARS-CoV-2-S_trunc_) were co-transfected with pHIV_NL_GagPol and pCCNanoLuc2AEGFP plasmids in HEK293T cells using FuGENE HD (Promega). Supernatant from transfected cells was harvested at 72 hr post transfection, filtered, stored at −80°C and subsequently titrated on HEK293T_ACE2_ cells.

### Pseudotyped virus neutralization assays

Neutralization assays were performed as previously (8). In brief, serially diluted monoclonal antibodies were incubated with the SARS-CoV-2 pseudotyped virus for 1 hour at 37°C degrees prior to adding to HEK293T/ACE2/TMPRSS2 mCherry cells (BEI Resources NR-55293) for 48 hours. Upon washing with PBS once, cells were lysed with Luciferase Cell Culture Lysis 5x reagent (Promega) and Nanoluc Luciferase activity of lysates measured using the Nano-Glo Luciferase Assay System with GloMax Discover System reader (Promega). Luminescence units were relative to those derived from cells infected with SARS-CoV-2 pseudotyped virus in the absence of monoclonal antibodies. The half-maximal inhibitory concentration of monoclonal antibodies (IC_50_) was determined using four-parameter nonlinear regression curve fit (GraphPad Prism).

### Antibody Protein Production

Antibodies were produced either as IgGs or Fab fragments, with some Fabs also being produced from papain cleavage of expressed IgGs (see below). For both, heavy chain and light chains for each antibody were co-transfected in Expi293F cells following manufacturer instructions and using the Expi293 Expression System Kit (Gibco, cat. no. A14635) in a 1:1 ratio. Cell supernatants were harvested, filtered, and subsequently ran over HiTrap MabSelect SuRe columns (Cytiva) to capture expressed IgGs or HisTrap columns (Cytiva) to capture expressed Fabs. Further purification was performed via SEC on either a HiLoad 16/600 Superdex 200 pg or Superdex 200 Increase 10/300 GL column (Cytiva) using an AKTA pure M1 system (Cytiva).

Antibody Fabs were also prepared via papain cleavage of IgGs expressed, as described above. In brief, papain (Sigma-Aldrich, cat. no. P3125) was incubated for 10 min in 2x Digestion Buffer (50 mM NaPO4 pH 7.0, 20 mM ethylenediaminetetraacetic acid (EDTA)) at 37°C at a final mass 1.5% (w/w) of the total IgG protein to be cleaved. Activated papain was then added to the IgGs at a 1:1 ratio (v/v) and incubated at 37°C for 1 hr with shaking at ∼100 rpm. The enzymatic reaction was quenched with iodoacetamide (Sigma-Aldrich, cat. no. I1149) and then ran over a HiTrap MabSelect SuRe column (Cytiva) to separate uncleaved IgG and cleaved Fc from the desired Fab product. Cleaved Fabs were further purified via SEC on a HiLoad 16/600 Superdex 200 pg or Superdex 200 Increase 10/300 GL column (Cytiva) after confirmation of separated IgG reactants and Fab products via SDS-PAGE gel.

### Antigen Protein Production

Sequences for the recombinant spike proteins (including RBDs) were codon optimized and cloned into a mammalian expression vector with an N-terminal secretory signal peptide and a variation of C-terminal tags including a the T4 Foldon trimerization domain, polyhistidine tag for affinity purification, and biotinylation motif. Plasmids were transfected in Expi293F following manufacturer instructions and using the Expi293 Expression System Kit (Gibco, cat. no. A14635). To obtain biotinylated proteins, Expi293F cells were co-transfected with a 1:1 ratio of the spike construct DNA and a plasmid encoding for the *Escherichia coli* biotin ligase (BirA) enzyme. Recombinant proteins were subsequently isolated from cellular supernatants via affinity chromatography using a HisTrap column (Cytiva). Further purification was performed via SEC on either a HiLoad 16/600 Superose 6 pg or Superose 6 Increase 10/300 column (Cytiva) using an AKTA pure M1 system (Cytiva).

### Enzyme-Linked Immunosorbent Assays (ELISAs)

ELISAs were used to measure the binding cross-reactivity of antibody IgGs to SARS-CoV-2 variant RBD proteins. High-binding 96-well assay plates were coated overnight at 4°C using solutions of 2 µg/mL RBD protein diluted in 1x TBS-Az. They were then washed with TBST-T (1x TBS + 0.05% Tween-20), blocked for 1 hr at room temperature in TBS-TMS (TBS-T + 1% non-fat dry milk + 1% goat serum), and then washed and coated with 10-fold serial dilutions in TBS-TMS with a top concentration of 10 µg/mL of monoclonal antibody IgGs for 2 hr. The plates were then washed with TBS-T and HRP-conjugated goat anti-human Ig Fc (Southern Biotech, cat. no. 2047-050) secondary antibody was added at a 1:4000 final dilution in TBS-TMS. Plates were washed again and then read out with 1-Step Ultra TMB-ELISA substrate (Thermo Scientific, cat. no. 34029), quenched with 1 M HCl solution, and absorbance was measured at 450 nm with a Tecan Infinity M Plex microplate reader using iControl 2.0 software. EC_50_ were calculated using the [Agonist] vs. response ---Variable slop (four parameters) model in GraphPad Prism v10.

### Biolayer Interferometry (BLI)

BLI assays were performed on an Octet RED96 system (FortéBio, Sartorius) at room temperature (20-25°C) in Octet Buffer (1x TBS + 0.1% bovine serum albumin + 0.02% Tween-20). For assays used to calculate binding kinetics, biotinylated RBD or Spike proteins were immobilized on streptavidin (SA) biosensors (Sartorius, cat. no. 18-5019) and dipped into a concentration series of monomeric Fabs. After subtraction of reference signal, the *k*_a_ (on rate), *k*_d_ (off rate), and dissociation constant (K_d_) were derived using the Association then Dissociation module in GraphPad Prism v10. For the ACE2 competition assay, biotinylated Spike protein was immobilized on SA biosensors and dipped into 2 µM antibody IgG for 300 seconds, followed immediately by association in 1 µM s-HACE2 for another 300 seconds.

### RBD Sequence Conservation Analysis

The ConSurf server (25) was used to autogenerate a selection of 48 CoV sequences that was aligned and assessed for sequence conservation based on a reference sequence and model for Wuhan-Hu-1 RBD (7RKU). Outputs of the server were used for conservation calculations as well as model coloring in Fig. 4.

### Evaluation of sarbecovirus cross-reactivity via high-throughput yeast-display binding assays

The complete pipeline for measuring mAb breadth across the pan-sarbecovirus panel is described at: https://github.com/tstarrlab/SARSr-CoV_mAb-breadth_Barnes/blob/main/results/summary/summary.md.

MAb binding via high-throughput FACS-seq was evaluated against a previously published pan-sarbecovirus panel of yeast-displayed RBDs (26) in the AWY101 yeast strain (27).

The yeast-display RBD library was grown, induced for yeast-surface expression, and labeled with monoclonal antibody at 10,000, 400, 16, 0.64, 0.0256, and 0 ng/mL concentration for one hour at room temperature. Yeast were washed with PBS-BSA and labeled with secondary FITC-conjugated chicken anti-Myc antibody (Immunology Consultants CMYC-45F) and PE-conjugated goat anti-human-IgG (Jackson ImmunoResearch 109-115-098). Libraries were then partitioned at each labeling concentration into four bins of mAb binding on a BD FACSAria, collecting a minimum of 1 million RBD^+^ cells per sample concentration across the four bins. Bins were set such that in a positive-binding control antibody:RBD combination (S2H97:WH1), 95% of cells would be sorted into bin 4 (Figure S2A). Conversely, in a negative-binding control antibody:RBD combination (no Ab:WH1), bins were set such that 95% of cells would be collected in bin 1 (Figure S2A). The remaining bin-boundary was set to bisect the distance on the PE-signal axis between bins 1 and 4. Cells were grown post-sort, plasmid purified, N16 barcode amplified, and sequenced on an Illumina NextSeq. Barcode reads were mapped to library barcodes, with raw counts of each barcode in each FACS bin found at: https://github.com/tstarrlab/SARSr-CoV_mAb-breadth_Barnes/blob/main/results/counts/variant_counts.csv.

For each library barcode, an EC50 binding strength was derived from its distribution of sequence reads across sort bins. First, the strength of mAb binding to each barcode at each mAb dilution was determined as the simple mean bin from cell counts across integer-weighted bins. Any barcode with less than 2 cell counts at any single sample concentration or less than an average of 5 cell counts across all sample concentrations was eliminated from analysis. An EC50 metric was then calculated from the fit of a sigmoidal curve between mean bin (mAb binding) and mAb labeling concentration. Per-barcode EC50 calculation and representative titration curve-fits can be found at: https://github.com/tstarrlab/SARSr-CoV_mAb-breadth_Barnes/blob/main/results/summary/compute_EC50.md, and per-barcode EC50 metrics are available at: https://github.com/tstarrlab/SARSr-CoV_mAb-breadth_Barnes/blob/main/results/bc_mAb_EC50/bc_mAb_EC50.csv.

We then computed the per-variant EC50 as the robust mean of replicate barcodes linked with the identical RBD variant, by taking the mean per-barcode EC50 after trimming tails of the top and bottom 5% of EC50 values among the replicate barcodes. The final variant derivation can be found at: https://github.com/tstarrlab/SARSr-CoV_mAb-breadth_Barnes/blob/main/results/summary/collapse_barcodes_lib61_SARSr-wts.md, and final per-variant mAb-binding values are available at: https://github.com/tstarrlab/SARSr-CoV_mAb-breadth_Barnes/blob/main/results/final_variant_scores/final_variant_scores_lib61.csv

### Evaluation of escape mutants via yeast-display deep mutational scanning

Deep mutational scanning libraries for SARS-CoV-2 variants Wuhan-Hu-1, Omicron XBB.1.5, and Omicron BA.2.86 were described in prior publications, including library construction, library availability, and measurements of mutational impacts on RBD expression and ACE2-binding affinity (28). These libraries consist of virtually all single amino acid changes in each RBD background in a yeast-surface display platform.

Duplicate yeast-display deep mutational scanning libraries were induced for RBD expression, and 5 OD^∗^mL of yeast were incubated in 1 mL for one hour at room temperature with a concentration of mAb corresponding to the EC90 of the mAb for the respective yeast-displayed wildtype RBD determined from pilot isogenic binding assays. In parallel, for FACS gate setting, 0.5 OD^∗^mL of the respective wildtype parental constructs were incubated in 100 μL of antibody at the matched EC90 concentration or 1/10 the EC90 concentration. Cells were washed, incubated with 1:100 FITC-conjugated chicken anti-Myc antibody to label RBD expression and 1:200 PE-conjugated goat anti-human-IgG to label bound antibody, and washed in preparation for FACS.

Antibody-escape cells in each library were selected via FACS on a BD FACSAria II or Cytek Aurora Cell Sorter. FACS selection gates were drawn to capture approximately 50% of yeast expressing the parental RBD control labeled at the 10x reduced antibody labeling concentration (see representative gating scheme in Fig. S2B). For each sample, 4 million RBD+ cells were processed on the sorter with collection of cells in the antibody-escape bin, which were expanded overnight, plasmid purified, and barcodes sequenced on an Illumina NextSeq. In parallel, plasmid samples were purified from 30 OD^∗^mL of pre-sorted library culture and sequenced to establish pre-selection barcode frequencies.

Demultiplexed Illumina barcode reads were matched to library barcodes and associated RBD mutant from previously assembled barcode-variant lookup tables using dms_variants (version 0.8.9), yielding a table of counts of each barcode in each pre- and post-sort population, available at: https://github.com/tstarrlab/SARS-CoV-2-RBD_Omicron_MAP_Barnes/tree/main/results/counts.

The escape fraction of each barcoded variant was computed from sequencing counts in the pre-sort and antibody-escape populations via the formula:

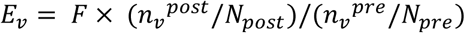

where *F* is the total fraction of the library that escapes antibody binding (e.g. annotated numbers in ‘escape’ gate, Fig. S2B), *n_v_*is the counts of variant *v* in the pre- or post-sort samples with a pseudocount addition of 0.5, and *N* is the total sequencing count across all variants pre- or post-sort. These escape fractions represent the estimated fraction of cells expressing a particular variant that fall in the escape bin, which scales from 0 for a mutation that never causes sufficient loss of binding to drive cells into the antibody-escape bin, to 1 for a mutation that escapes binding >10-fold such that the variant falls into the antibody-escape bin defined by the control labeling gates. We applied computational filters to remove mutants with low pre-selection sequencing counts or highly deleterious mutations that escape antibody binding artefactually due to poor RBD surface expression, specifically mutants with orthogonally measured ACE2-binding impacts of <–3 (1000-fold loss of ACE2 binding) or expression scores of <–1.25, <–1.25, and <–0.75 for Wuhan-Hu-1, XBB.1.5, and BA.2.86, respectively, accounting for the variation in baseline expression levels of different wildtype variants and differences in the arbitrary units scaling of expression mean fluorescence intensity across experiments. Final per-mutant escape fractions were computed as the average across barcodes within replicates, with the correlation between replicate library selections shown in Fig. S2C. Final escape fraction measurements averaged across replicates are available at: https://github.com/tstarrlab/SARS-CoV-2-RBD_Omicron_MAP_Barnes/tree/main/results/supp_data.

### Mouse challenge experiments

Mouse challenge studies were performed using a previously described K18-hACE2 SARS-CoV-2 infection model with modifications for the antibody evaluated in this study (29). All animal procedures were approved by the Institutional Animal Care and Use Committee at The Rockefeller University (protocol 21042-H). Eight-week-old K18-hACE2 transgenic mice [B6.Cg-Tg(K18-ACE2)2Prlmn/J; Jackson Laboratory, strain 034860] were maintained under specific pathogen-free ABSL3 conditions with unrestricted access to food and water. Mice (n = 5 per group) received C5078 by intraperitoneal injection at 5 mg kg⁻f either 24 h before intranasal challenge (prophylactic group) or 4 h after infection (therapeutic group). Animals were infected with 2 × 10⁴ PFU of SARS-CoV-2 XBB.1.5 by the intranasal route. At 3 days post-infection, the left lung was harvested and homogenized in TRIzol LS reagent (Ambion, Thermo Fisher Scientific). Total RNA was isolated by chloroform phase separation, precipitated with isopropanol, washed with 75% ethanol, and resuspended in nuclease-free water. Viral RNA levels were quantified by one-step quantitative RT-PCR using the Power SYBR Green RNA-to-CT 1-Step Kit (Thermo Fisher Scientific) on a StepOnePlus Real-Time PCR System (Applied Biosystems). Amplification targeted the SARS-CoV-2 nucleocapsid (N) gene using primers 2019-nCoV_N1-F (5′-GACCCCAAAATCAGCGAAAT-3′) and 2019-nCoV_N1-R (5′-TCTGGTTACTGCCAGTTGAATCTG-3′). Quantification was performed using the 2019-nCoV_N Positive Control (IDT) as a standard.

### Cryo-EM Sample Preparation

For the C5078–RBD–S309–S2X259 complex, purified RBD (res 328-533) and Fabs were combined at a molar ratio of 2:2:2:1 C5078 Fab:S309 Fab:S2X259 Fab:RBD and incubated at room temperature overnight. Following SEC on a Superdex 200 Increase 10/300 GL column (Cytiva), a fraction from the complex-containing peak was isolated and concentrated to 10 mg/mL for grid preparation. UltrAufoil Au R0.6/1, 300 mesh grids (Electron Microscopy Sciences) were glow-discharged for 90 s at 15 mA on an easiGlow (PELCO) before sample application. Immediately prior to depositing 3.1 uL of the complex onto the glow-discharged grid, fluorinated octyl-maltoside (Anatrace) was added to the sample to a final concentration of 0.002% w/v. Using a Vitrobot Mark IV (Thermo Fisher Scientific), the grid was blotted for 3 s at room temperature, 100% humidity, and with blot force 3. It was then plunge-frozen in liquid ethane and transferred to liquid nitrogen for storage.

For B2014 and C5078 Fab – SARS-CoV-2 BA.1 S6P complexes, freshly purified SARS-CoV-2 BA.1 S6P was incubated with 1.1 molar excess of cleaved Fab for 10-20 min in 1x TBS. Quantifoil Cu R1.2/1.3 300 mesh grids (Electron Microscopy Sciences) were glow-discharged for 60 s at 10 mA on an easiGlow (PELCO) before sample application. Immediately prior to depositing 3.1 uL of the 3.2-3.5 mg/mL complex onto the glow-discharged grids, fluorinated octyl-maltoside (Anatrace) was added to the samples to a final concentration of 0.025% w/v. Using a Vitrobot Mark IV (Thermo Fisher Scientific), grids were blotted for 3-4 s at room temperature, 100% humidity, and with blot force 0 or 4. They were then plunge-frozen in liquid ethane and transferred to liquid nitrogen for storage.

### Cryo-EM Data Collection and Processing

#### High Resolution C5078–RBD–S309–S2X259 and B2014–BA.1 S6P Cryo-EM Data Collection & Processing

Movies were collected on a 300 kV TFS Krios microscope with a Falcon4i camera at 130kx magnification and 0.92 Å/pix using EPU software. They were then imported into cryoSPARC v4.1 (30) for all processing steps: beginning with motion correction, CTF estimation, micrograph curation, blob picking, and particle extraction. Particles were first extracted at a box size that had been Fourier cropped to half its original size for initial processing, followed by 2D classification and then 3D classification (using tandem ab-initio and heterogeneous refinement jobs) before re-extraction at the full box size. For the C5078–RBD–S309– S2X259 dataset, a small subset of micrographs was first extracted at full box size and subjected to ab-initio reconstruction to populate volumes for heterogeneous refinement of the full dataset. Additionally, the B2014–BA.1 S6P dataset underwent another round of heterogenous refinement at the full box size. Non-uniform refinement was then used to generate global maps for each complex and then local refinement with masks for regions of interest applied generated the local maps with better features at the antibody-antigen interfaces. Further details of data collection and processing workflows are provided in Fig. S4 and Table S1.

#### Low Resolution C5078–BA.1 S6P Cryo-EM Data Collection & Processing

Movies were collected on a 200 kV TFS Glacios microscope with a Falcon4i camera at 120kx magnification and a pixel size of 1.17 Å/pix using EPU software. Movies were imported into cryoSPARC for all processing steps: beginning with motion correction, CTF estimation, micrograph curation, blob picking, and particle extraction. Particles were first extracted at a 432 pix box size that had been Fourier cropped to 216 pix for initial processing. One round each of 2D and 3D classification (using tandem ab initio and heterogeneous refinement jobs) preceded re-extraction at the full box size. From here, further 2D and 3D classification was carried out, followed by global CTF refinement of the final particle stack. Non-Uniform refinement with C1 symmetry applied generated the final map. While the GSFSC reports 4.27 Å resolution, map features appear more consistent with a lower resolution map. More detail is provided in Table S1.

### Modeling and Analysis of cryo-EM structures

Initial model coordinates for each protein were obtained by docking individual chains from homology models, identified using IMGT/3Dstructure-DB (31) (Table S1), into the sharpened cryo-EM maps using UCSF ChimeraX (32). Sequences of the docked models were then updated manually in Coot (33) to align with the expressed protein. Finally, models were refined by using iterative rounds of refinement in Phenix (34) and manual building and refinement in Coot. UCSF ChimeraX was used to visualize structures, create figures, and define contacts using a 4.1 Å inter-chain distance cutoff. Buried surface area (BSA; sum of heavy and light chain interfaces) and potential hydrogen bond assignments described were calculated using a 1.4 Å probe in PDBePISA (35). Antibody residues were numbered using Kabat convention and CDRs defined using IMGT designation (36). Antibody germline sequence analyses were done using either IMGT V-Quest (37) or IgBlast (38) servers online.

## ACKNOWLEDGEMENTS

We thank members from the laboratory of Dr. Pamela Bjorkman (California Institute of Technology) and Dr. Peter S. Kim (Stanford) for cell lines, SARS-CoV-2 spike expression constructs, and technical assistance with pseudovirus neutralization assays. Cryo-EM data for this work was collected at the Stanford-SLAC cryo-EM resource center with support from Dr. Bharti Singal and Dr. Haoqing Wang. We thank the University of Utah Flow Cytometry Core and Center for High Throughput Computing facilities. P.D.B., and M.C.N. are Howard Hughes Medical Institute (HHMI) Investigators and C.O.B. is a HHMI Freeman Hrabowski Scholar. This article is subject to HHMI’s Open Access to Publications Policy. HHMI lab heads have previously granted a non-exclusive CC BY 4.0 license to the public and a sublicensable license to HHMI in their research articles. Pursuant to those licenses, the author-accepted manuscript of this article can be made freely available under a CC BY 4.0 license immediately upon publication. Finally, we thank all members of the Barnes lab for their careful review and feedback on the manuscript.

## AUTHOR CONTRIBUTIONS

MEA and COB conceptualized and designed the project. MEA and COB performed and analyzed structural biology data. MEA, JAL, and DV performed and analyzed binding experiments, with support from YEL and ME in protein expression and purification. WBF and TNS performed and analyzed deep mutational scanning data. VAB performed and analyzed mouse challenge experiments. PDB, MCN, TNS, and COB acquired funding and supervised the work. MEA, WBF, TNS and COB visualized the data. MEA and COB wrote the original draft and received review and editing from all authors.

## FUNDING

This study was supported in part by funds from the Howard Hughes Medical Institute Emerging Pathogens Initiative (C.O.B). Additionally, C.O.B. acknowledges support by the HHMI Hanna Gray Fellowship, Rita Allen Foundation, and Pew Biomedical Scholars Program. T.N.S acknowledges support from the Searle Scholars Program and NIH/NIAID (DP2 AI177890). V.A.B. is supported by the Boehringer Ingelheim Fonds PhD fellowship. Z.W. received support from the SNF Institute for Global Infectious Disease for the Advancement of Translational Research, partially funded by grant # UL1 TR001866 from the National Center for Advancing Translational Sciences (NCATS), under the National Institutes of Health (NIH) Clinical and Translational Science Award (CTSA) program. P.D.B, and M.C.N are Howard Hughes Medical Institute investigators. C.O.B. is a HHMI Freeman Hrabowski Scholar.

## CONFLICTS OF INTEREST

The Rockefeller University has filed a provisional patent application in connection with monoclonal antibodies described in this work on which Z.W. and M.C.N. are inventors (US patent 17/575,246).

## DATA AND CODE AVAILABILITY

The atomic models and cryo-EM maps generated for RBD C5078–RBD–S309–S2X259 (global refinement), C5078–RBD (local refinement), and B2014–BA.1 RBD (local refinement) have been deposited to the Protein Data Bank (PDB; http://www.rcsb.org/) and the Electron Microscopy Databank (EMDB; http://www.emdataresource.org/) under accession codes PDB: 37IZ, 37JA, and 37JB and EMD: −78223, −78224, and −78226 respectively. Cryo-EM maps generated for B2014–BA.1 S6P (global refinement) and C5078–BA.1 S6P (global refinement) have been deposited to the EMDB under accession codes −78225 and −78227, respectively. The code and pipeline used to analyze the pan-sarbecovirus breadth and deep mutational scanning data is available at, respectively, https://github.com/tstarrlab/SARSr-CoV_mAb-breadth_Barnes/tree/main and https://github.com/tstarrlab/SARS-CoV-2-RBD_Omicron_MAP_Barnes/tree/main.

## Supplementary Material

**Supplementary Figure 1.**
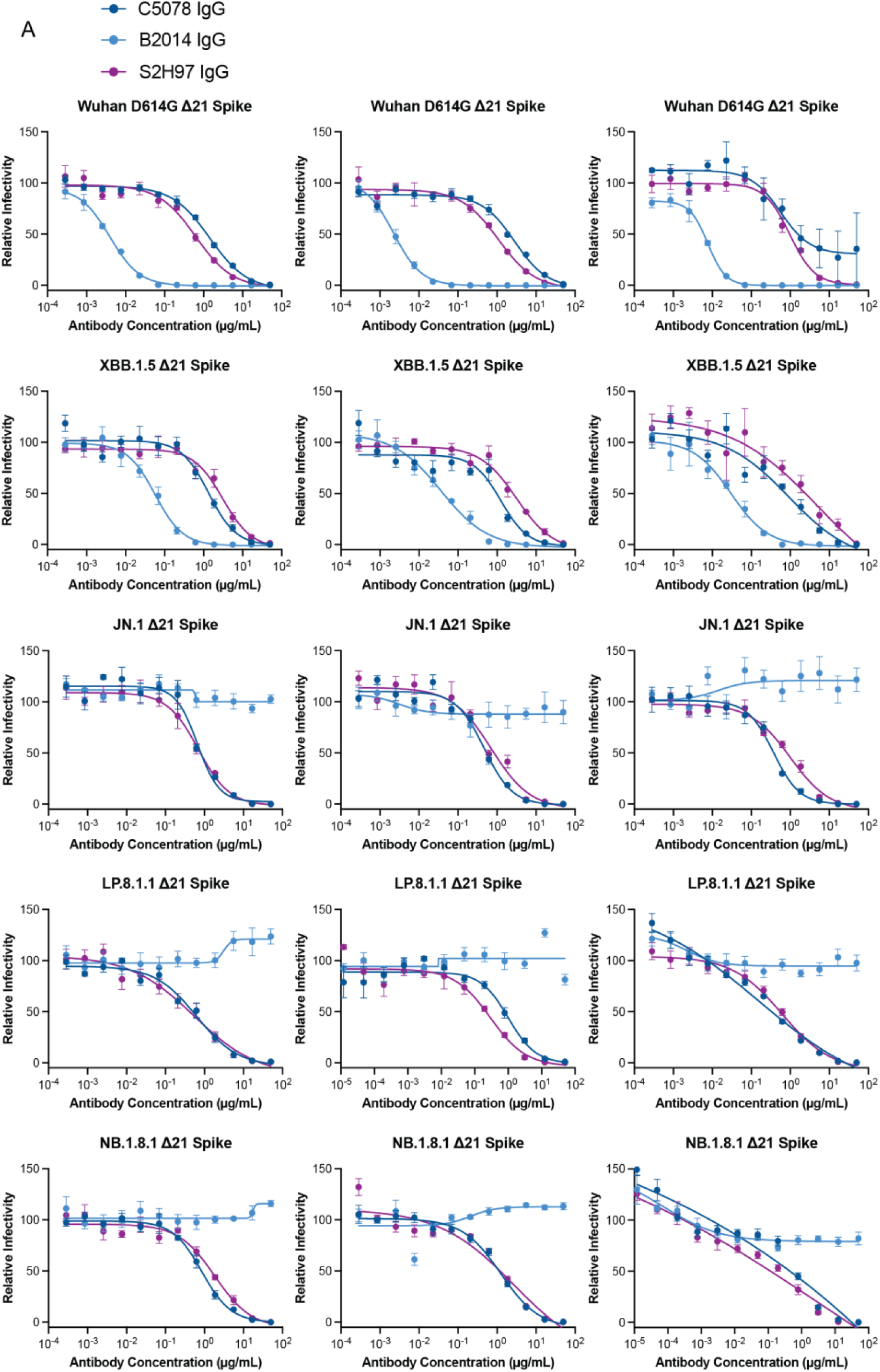
Pseudovirus neutralization against SARS-CoV-2 variants. **A)** Neutralization curves for three replicates each of SARS-CoV-2 pseudoviruses by C5078 IgG (dark blue), B2014 IgG (light blue), and S2H97 IgG (purple). Data points are the mean of quadruplicate values with standard error of the mean reported as error bars.

**Supplementary Figure 2.**
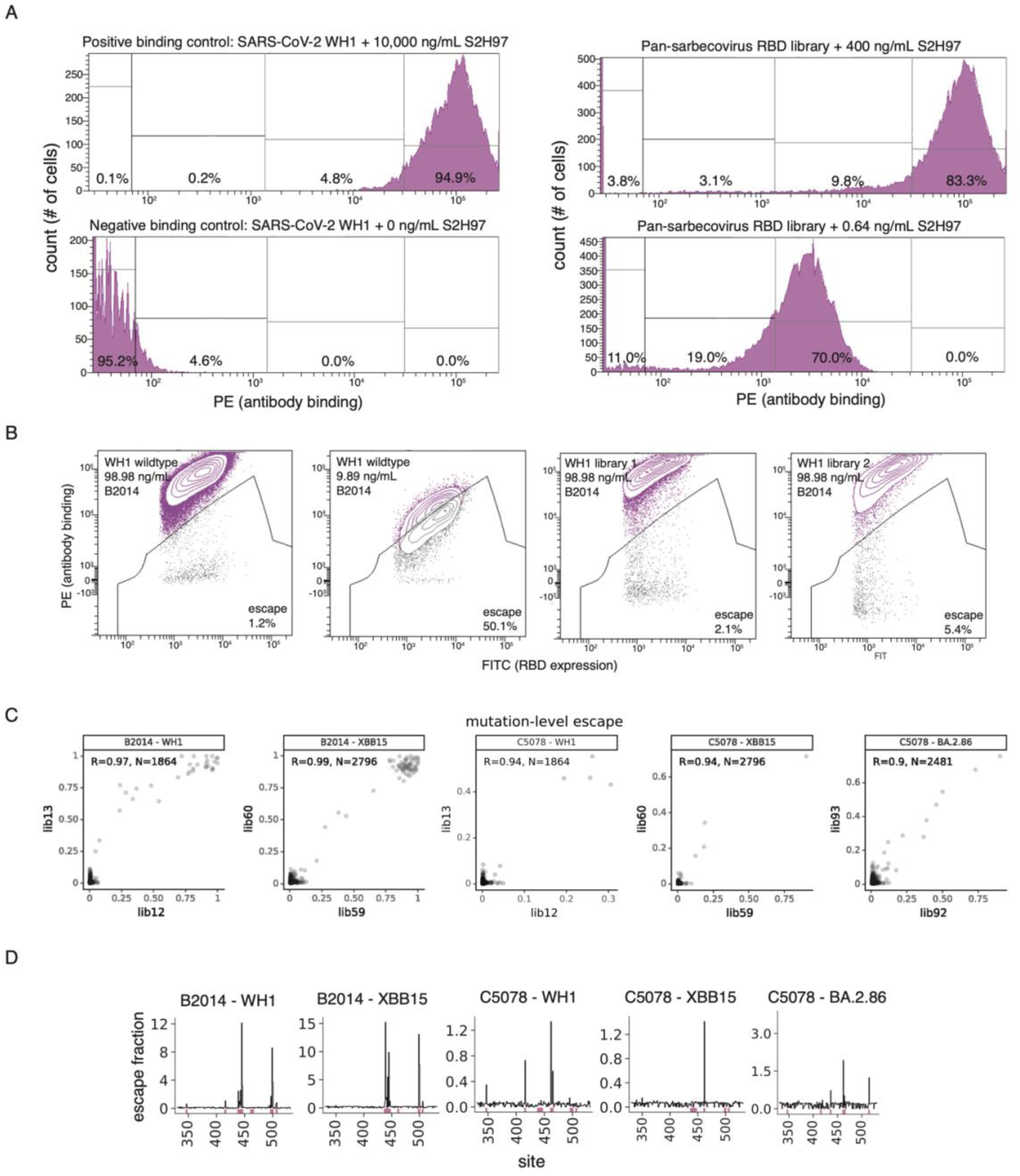
Yeast-display experimental design. **A)** Representative FACS gates used to identify breadth of B2014 binding amongst sarbecoviruses. Bin-boundaries 1:2 and 3:4 are drawn to capture 95% of negative-binding and positive-binding wildtype controls, respectively. Boundary 2:3 is drawn to bisect the remaining distance on the PE-signal axis. Percentages represent the fraction of the overall yeast population sorted into a given bin in each condition. **B)** Representative FACS gates used to identify mutations that escape B2014 binding. An antibody-escape gate was drawn to capture approximately 50% of the cells in the respective wildtype control labeled at 0.1x the library selection antibody concentration. “Escape fraction” represents the fraction of cells of a mutant genotype that fall into this antibody-escape FACS gate. **C)** Correlation in the per-mutant escape fraction of B2014 and C5078 across duplicate library selections. **D)** For each escape assay, lineplot shows the total escape of mutations at each site in the RBD. Sites of strong escape indicated by pink bars are visualized at the per-mutation level in logoplots (Figs. 2E, 4B).

**Supplementary Figure 3.**
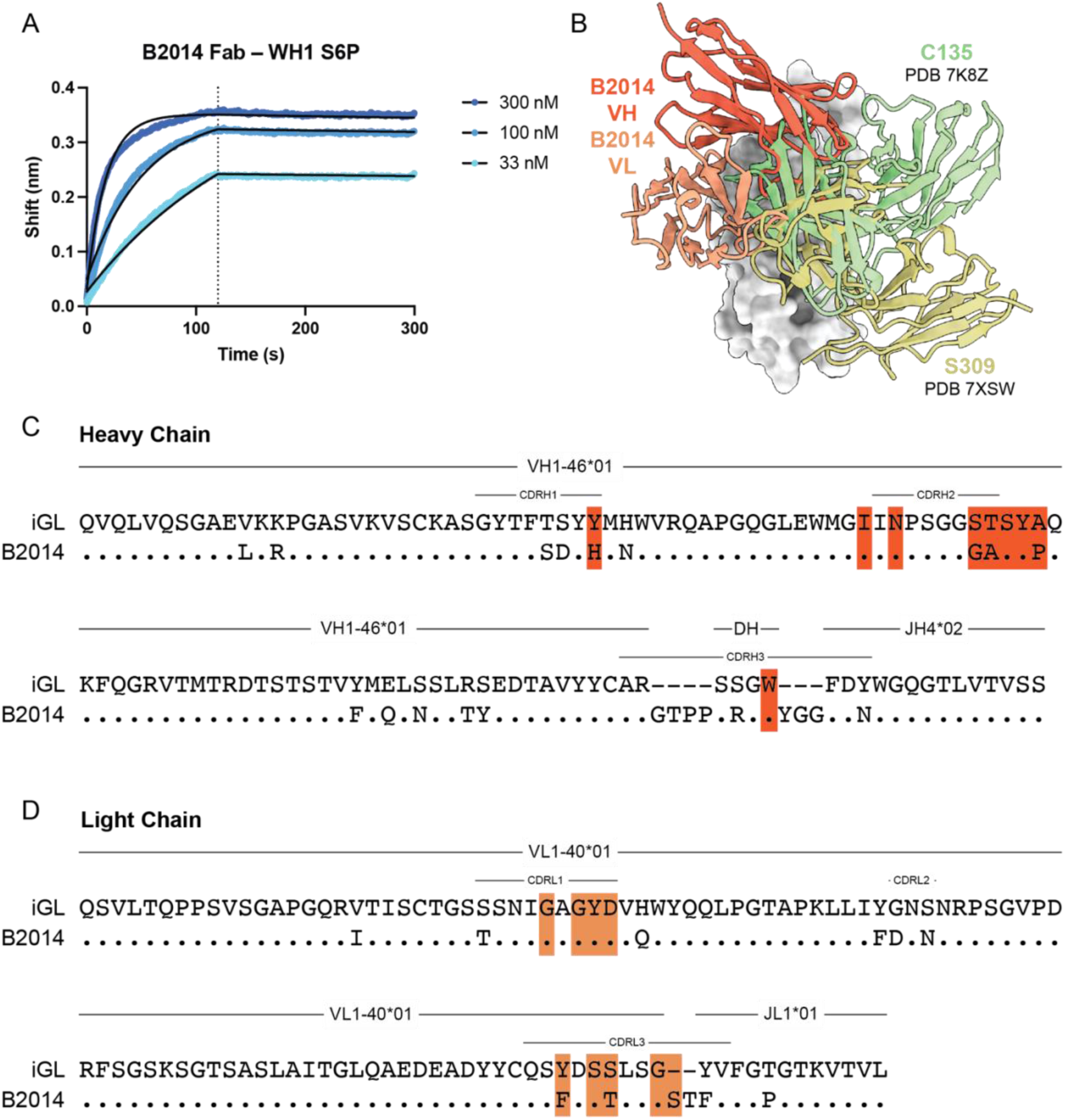
Characterization of B2014 molecular interactions with SARS-CoV-2 BA.1 Spike protein. **A)** BLI plot of the binding interactions between B2014 Fab and Wuhan-Hu-1 6P Spike at a range of concentrations. The dissociation phase begins at 120 seconds and is denoted by a dotted line. Fit curves are shown as black solid lines. **B)** RBD-VHVL model of B2014 shown with C135 and S309 RBD-VHVL models shown after alignment of the RBD subunits. **C,D)** Alignments of the B2014 **C)** Heavy chain and **D)** Light chain with their respective inferred germlines (iGLs), which were computed using IMGT V-Quest. Assigned germline genes and CDR loop definitions are shown above. Junction residues without germline assignments are represented with dashes. Contacts are identified with shaded orange boxes.

**Supplementary Figure 4.**
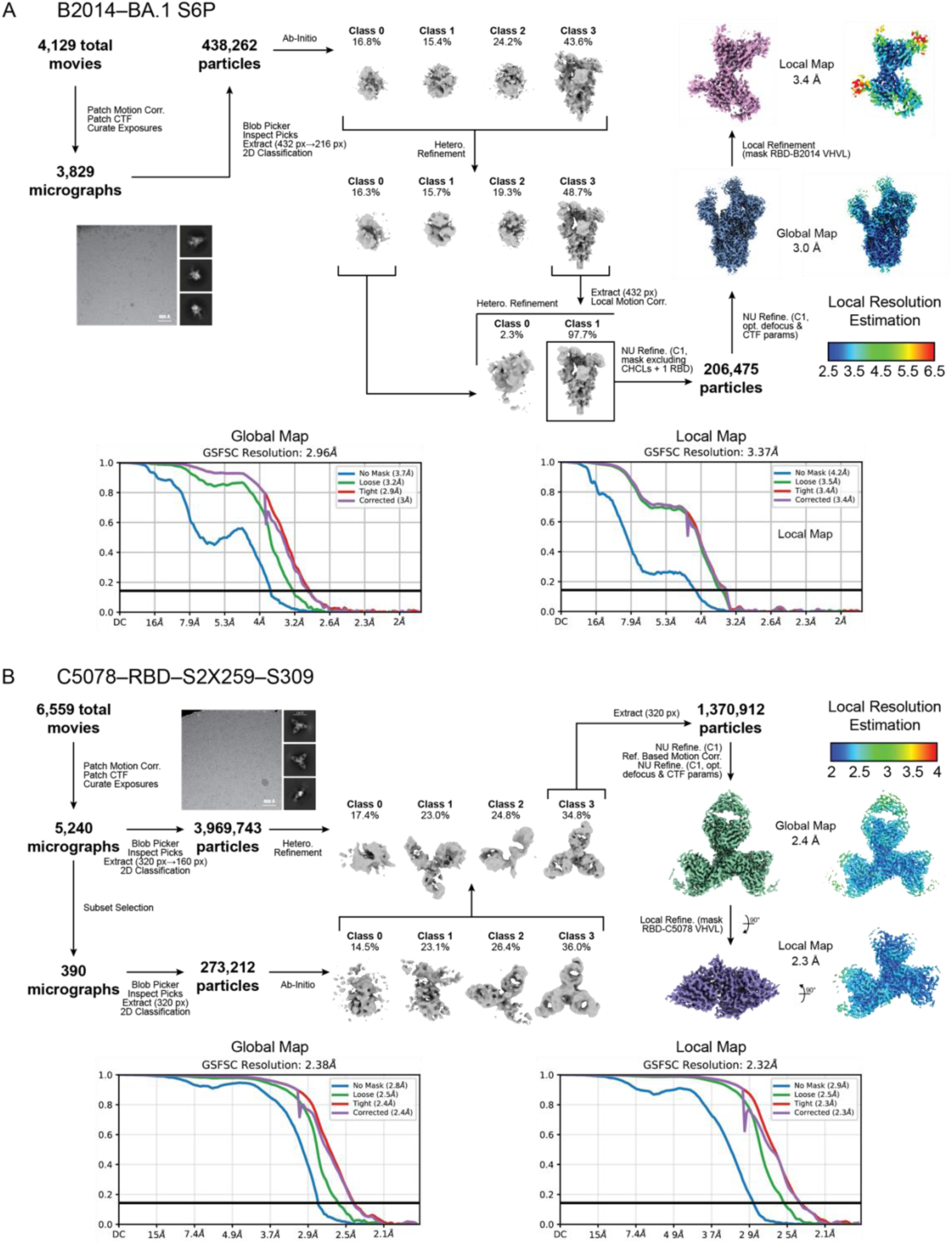
Cryo-EM data collection and processing workflows for B2014–BA.1 S6P and C5078–RBD– S2X259–S309. **A,B)** Cryo-EM data collection and processing workflows for **A)** B2014–BA.1 S6P and **B)** C5078–RBD– S2X259–S309 complex datasets, including processing details that resulted in the final global and local maps, as well as their local resolution estimation plots and Gold Standard FSC curves.

**Supplementary Figure 5.**
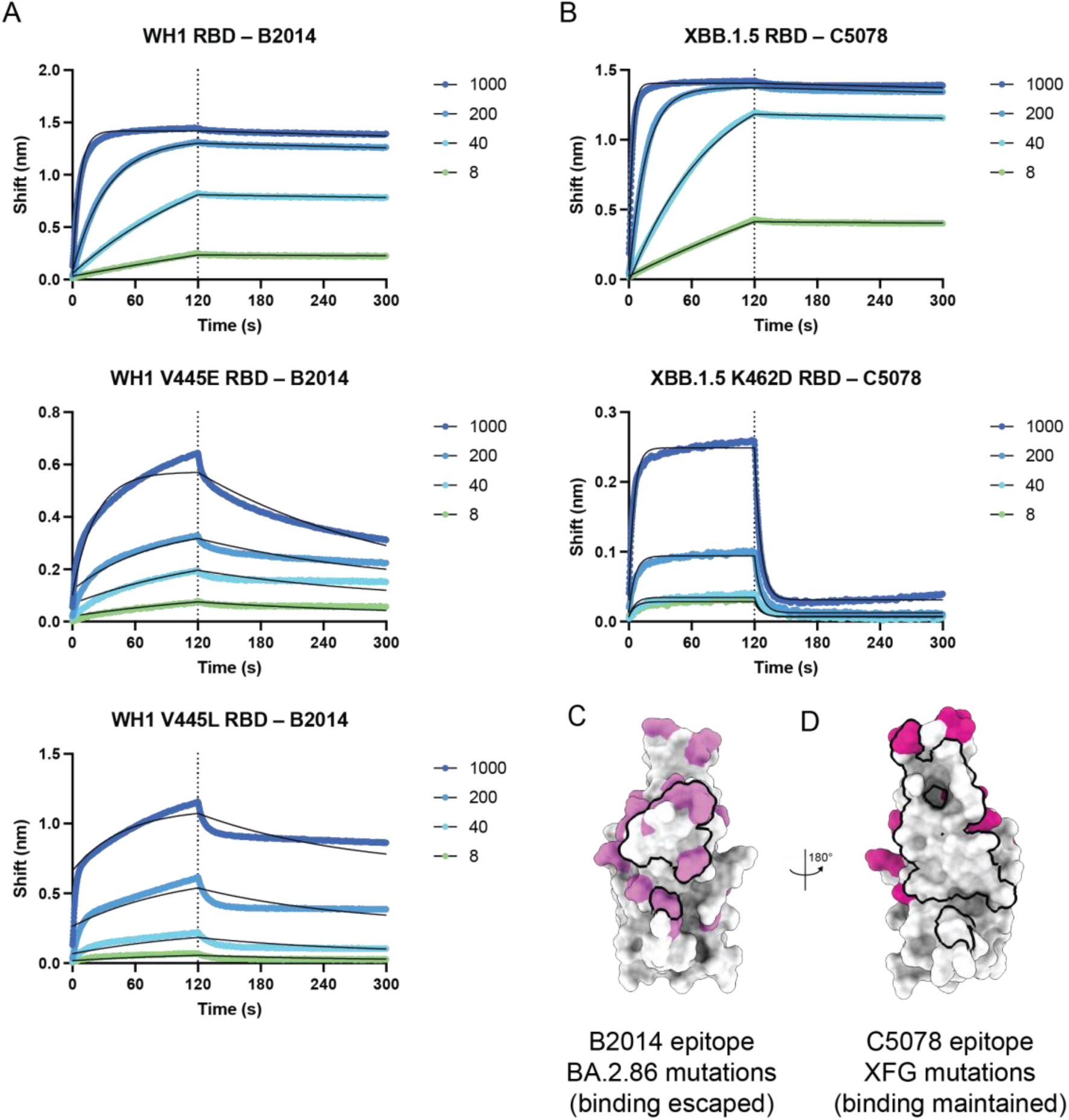
Validation of DMS-predicted escape mutations using biolayer interferometry (BLI). **A)** BLI plots of the binding interactions between B2014 IgG and WH1, WH1 V445E, and WH1 V445L RBDs. The WH1 RBD single point mutants were selected from the results of Fig. 2D. IgG was loaded onto ProA biosensors and then dipped into RBD in solution. Concentrations are in units of nM. **B)** BLI plots of the binding interaction between C5078 IgG and XBB.1.5 or XBB.1.5 K462D RBDs. The XBB.1.5 RBD single point mutant was selected from the results of Fig. 4B. IgG was loaded onto ProA biosensors and then dipped into RBD in solution. Concentrations are in units of nM. **C)** Model of the WH1 RBD (from this work) with residues included in the B2014 epitope outlined in black and residues mutated in the BA.2.86 strain (compared to Wuhan-Hu-1) colored in violet. Note from Fig. 1B that B2014 does not bind BA.2.86 RBD. **D)** Model of the WH1 RBD (from this work) with residues included in the C5078 epitope outlined in black and residues mutated in the XFG strain (compared to Wuhan-Hu-1) colored in magenta. Note from Fig. 1B that C5078 does bind XFG RBD.

**Supplementary Figure 6.**
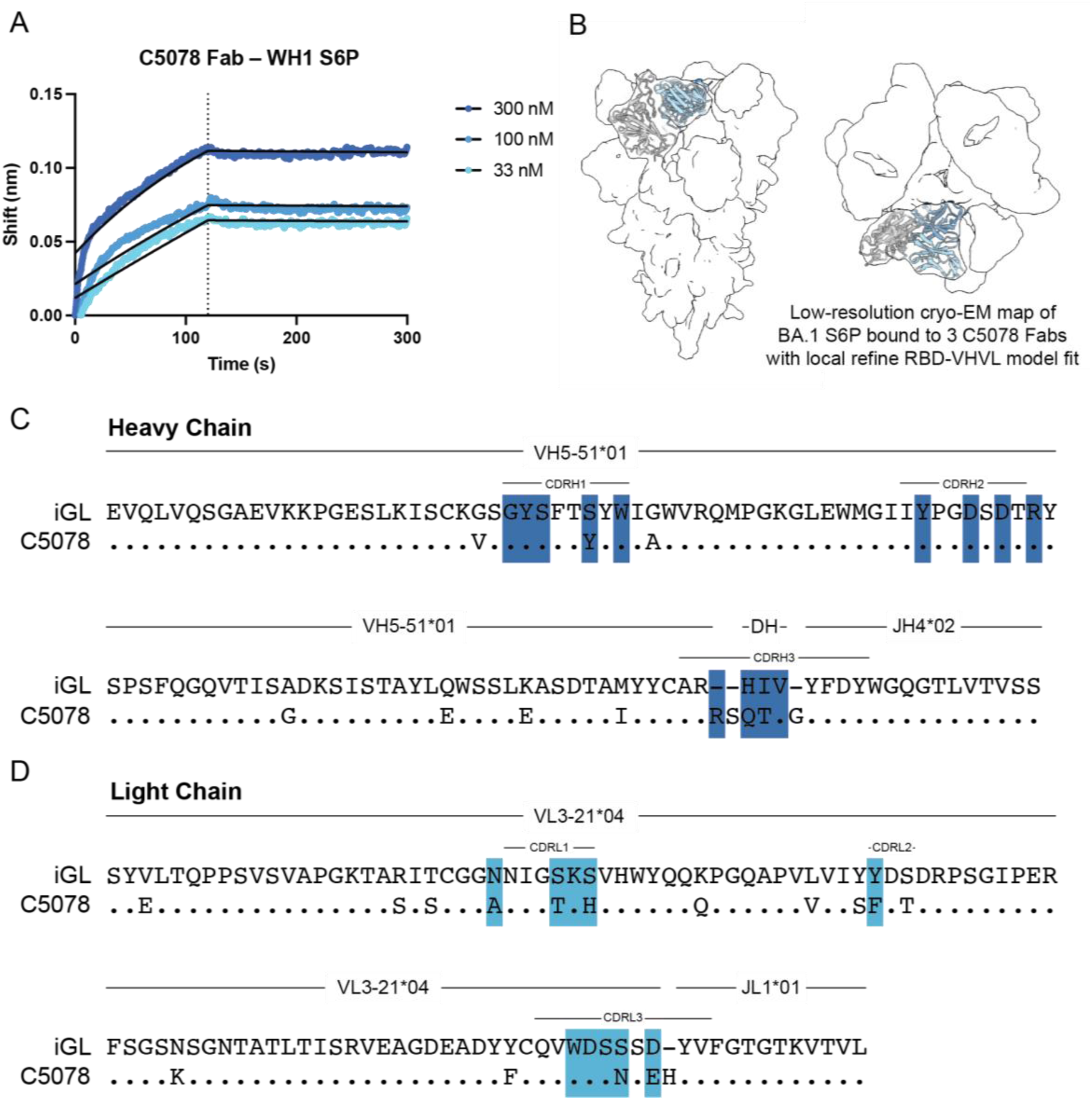
Characterization of C5078 molecular interactions with SARS-CoV-2 Spike protein. **A)** BLI plot of the binding interactions between C5078 Fab and Wuhan-Hu-1 6P Spike at a range of concentrations. The dissociation phase begins at 120 seconds and is denoted by a dotted line. Fit curves are shown as black solid lines. **B)** Low-resolution cryo-EM map of BA.1 S6P bound to 3 C5078 Fabs with local refine RBD-VHVL model (PDB 37JA) fit. **C,D)** Alignments of the C5078 D) Heavy chain and E) Light chain with their respective inferred germlines (iGLs), which were computed using IMGT V-Quest. Assigned germline genes and CDR loop definitions are shown above. Junction residues without germline assignments are represented with dashes. Contacts are identified with shaded blue boxes.

**Supplementary Figure 7.**
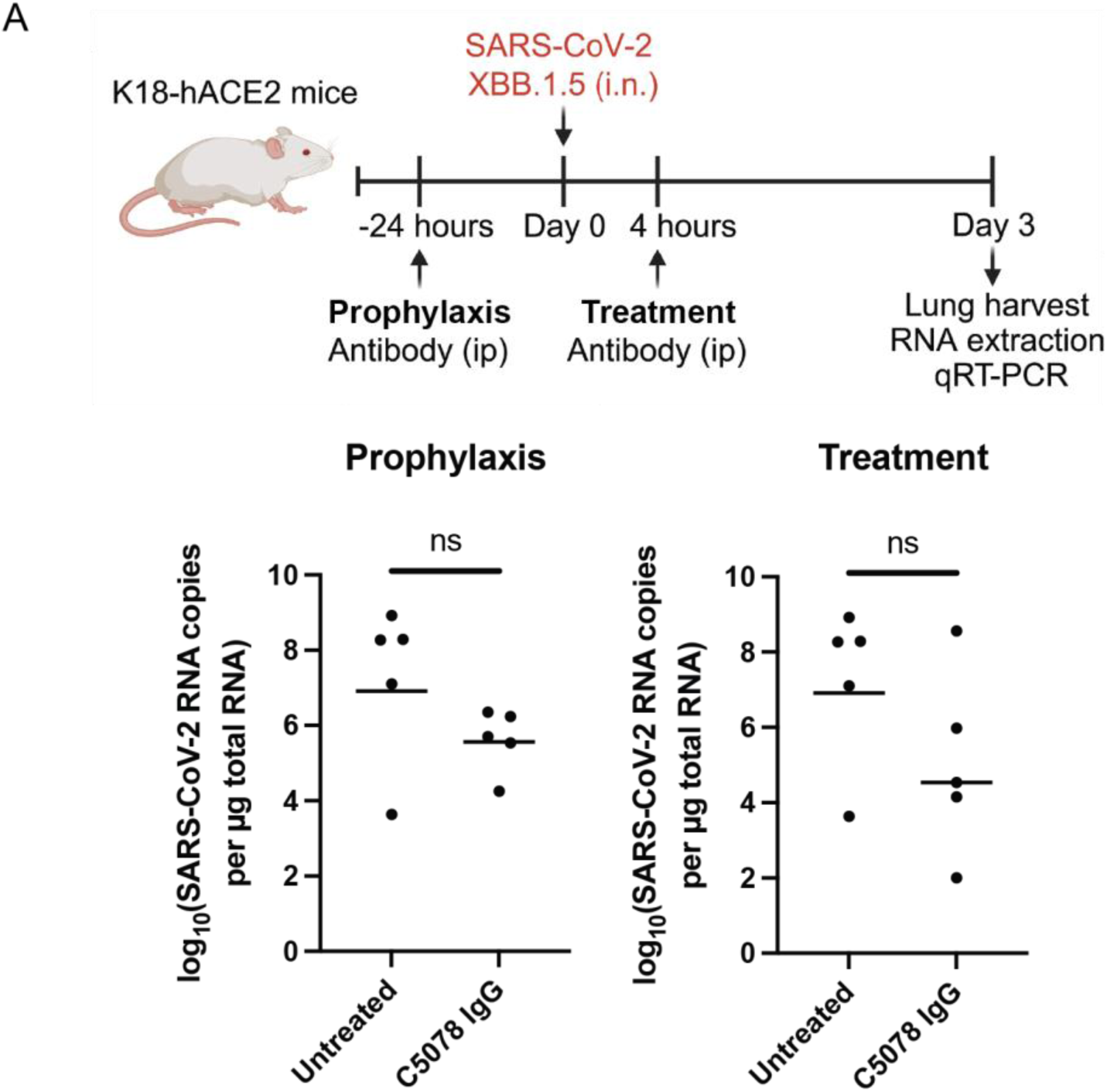
*In vivo* evaluation of C5078 against SARS-CoV-2 XBB.1.5. **A)** Schematic representation of *in vivo* experimental timeline followed by plots of the results for animals injected with C5078 IgG 24 hours prior to infection (Prophylaxis, left) or 4 hours after infection (Treatment, right). Total of n=5 mice per group. Data were analyzed using one-way ANOVA, correcting for multiple comparisons with Dunnett’s test with comparison to the untreated control. ns, not significant.

**Supplementary Table 1.** Cryo-EM data collection and processing statistics.

|  | C5078-RBD-S309-S2X259 |  | B2014-BA.1 S6P |  | C5078-BA.1 S6P |
| --- | --- | --- | --- | --- | --- |
|  | (global) | (local) | (global) | (local) | (global) |
| **PDB** | **37IZ** | **37JA** | n/a | **37JB** | n/a |
| **EMD** | **-78223** | **-78224** | **-78225** | **-78226** | **-78227** |
| **Data collection conditions** |  |  |  |  |  |
| Microscope | TFS Krios |  | TFS Krios |  | TFS Glacios |
| Camera | Falcon 4i |  | Falcon 4i |  | Falcon 4i |
| Magnification | 130,000x |  | 130,000x |  | 120,000x |
| Voltage (kV) | 300 |  | 300 |  | 200 |
| Dose rate (e<sup>-</sup>/pixel/s) | 6.7 |  | 11.9 |  | 9.8 |
| Electron dose (e<sup>-</sup>/Å<sup>2</sup>) | 50 |  | 50 |  | 30 |
| Defocus range (μm) | -0.8 to -2 |  | -1 to -2.5 |  | -1 to -2 |
| Pixel size (Å) | 0.92 |  | 0.92 |  | 1.17 |
| Micrographs collected | 6,559 |  | 4,129 |  | 6,491 |
| Micrographs used | 5,240 |  | 3,829 |  | 4,590 |
| Total extracted particles | 3,969,743 |  | 438,262 |  | 1,779,711 |
| Final refined particles | 1,370,912 |  | 206,475 |  | 267,805 |
| Symmetry imposed | C1 | C1 | C1 | C1 | C1 |
| Nominal Map Resolution (Å) |  |  |  |  |  |
| FSC 0.143 (unmasked/masked) | 2.8/2.4 | 2.9/2.3 | 3.7/3.0 | 4.2/3.4 | 5.2/4.3 |
| **Refinement and Validation** |  |  |  |  |  |
| Initial model | 8HN7, 7U0K, 7N3E, 7R6W, 7M7W |  | n/a | 7TN0, 7JXE, 5AZE | n/a |
| Model Resolution (Å) |  |  |  |  |  |
| FSC 0.143 | 2.4 | 2.3 | n/a | 3.3 | n/a |
| Number of atoms |  |  |  |  |  |
| Protein | 6,843 | 3,285 | n/a | 3,311 | n/a |
| Ligand | 0 | 0 | n/a | 0 | n/a |
| MapCC (global/local) |  |  |  |  |  |
| Map sharpening B-factor | -96.9 | -88.1 |  |  | -151 |
| R.m.s. deviations |  |  |  |  |  |
| Bond lengths (Å) | 0.005 | 0.004 | n/a | 0.004 | n/a |
| Bond angles (°) | 1.071 | 0.965 | n/a | 0.995 | n/a |
| MolProbity score | 1.65 | 1.27 | n/a | 1.87 | n/a |
| Clashscore (all atom) | 8.37 | 5.15 | n/a | 9.77 | n/a |
| Poor rotamers (%) | 1.08 | 0 | n/a | 0 | n/a |
| Ramachandran plot |  |  |  |  |  |
| Favored (%) | 97.04 | 98.31 | n/a | 94.84 | n/a |
| Allowed (%) | 2.85 | 1.69 | n/a | 5.16 | n/a |
| Disallowed (%) | 0.11 | 0 | n/a | 0 | n/a |

